# Paracrine factors of stressed peripheral blood mononuclear cells activate pro-angiogenic and anti-proteolytic processes in whole blood cells and protect the endothelial barrier

**DOI:** 10.1101/2022.04.22.489132

**Authors:** Dragan Copic, Martin Direder, Klaudia Schossleitner, Maria Laggner, Katharina Klas, Daniel Bormann, Hendrik Jan Ankersmit, Michael Mildner

## Abstract

Tissue regenerative properties have been attributed to secreted paracrine factors derived from stem cells and other cell types. Especially, the secretome of γ-irradiated peripheral blood mononuclear cells (PBMCsec) has been shown to possess high tissue-regenerative and pro-angiogenic capacities in a variety of preclinical studies. In the light of future therapeutic intravenous applications of PBMCsec, we investigated possible effects of PBMCsec on circulating white blood cells and endothelial cells lining the vasculature.

**Methods:** To identify changes in the transcriptional profile of white blood cells treated with PBMCSec, whole blood was drawn from healthy individuals and stimulated with PBMCsec for 8 hours *ex vivo* before further processing for single cell RNA sequencing (scRNAseq). In addition, we performed in vitro assay to confirm findings arising from the transcriptional profiling.

**Results:** Addition of PBMCsec to whole blood significantly altered the gene signature of granulocytes (17 genes), T-cells (45 genes), B-cells (72 genes) and most prominently monocytes (322 genes). We detected a strong upregulation of several tissue-regenerative and pro-angiogenic cyto- and chemokines in monocytes, including *VEGFA, CXCL1* and *CXCL5*. Intriguingly, inhibitors of endopeptidase activity, such as *SERPINB2*, were also strongly induced. Measurement of the trans-endothelial electrical resistance of primary human microvascular endothelial cells revealed a strong barrier-protective effect of PBMCsec after barrier disruption.

**Conclusion:** Together, we show that PBMCsec induces angiogenic and proteolytic processes in the blood and is able to attenuate endothelial barrier damage. These regenerative properties suggest that systemic application of PBMCsec might be a promising novel strategy to restore damaged organs.

## Introduction

Central goals in modern regenerative medicine are the repair of injured tissues and organs along with restoration of their innate functions [1]. Cell-based therapies have made tangible progress to this field over the past decades [2]. Mesenchymal stem cells (MSC) possess high capacities for self-renewal and differentiation which makes them a highly attractive option to promote tissue regeneration. However, in spite of encouraging pre-clinical data, most of the first-in-men clinical trials failed to show effectivity in patients [3][4]. Beyond that, it became increasingly apparent that released paracrine factors, rather than engraftment and differentiation of applied stem cells, are crucial for most of the beneficial biological effects and thus predominantly contribute to the observed tissue-regenerative effects [5–8]. In addition, stem cell secretome-based therapies bear considerable limitations, including the need for invasive procedures for isolation, low abundance and high costs for expansion and preservation. This highlights the need for alternative sources [9].

Recently, several studies suggested peripheral blood mononuclear cells (PBMCs) as a valuable alternative source to MSCs. In addition, γ-irradiation of PBMCs have been shown to promote the production and release of soluble factors and to exert tissue-protection [10]. In-depth functional analyses of the secretome of γ-irradiated PBMCs (PBMCsec) revealed a vast array of proteins, lipids and extracellular vesicles as the main biological constituents and confirmed that the presence of all fraction is required in order for PBMCsec to exert its biological effects to its full capacity [11][12]. The modes of action by which PBMCsec exerts its beneficial effects range from promotion of angiogenesis [11], antimicrobial activity [13], cytoprotection [14], immunomodulation [15], improvement of re-epithelization [16] to vasodilation and the inhibition of platelet aggregation [17]. More recently, Laggner and colleagues were able to demonstrate a decrease of dendritic cell-mediated skin inflammation in a murine model of contact hypersensitivity after topical treatment with PBMCsec, further expanding the broad spectrum of possible clinical implications of this investigational medicinal product [18]. Beneficial effects of PBMCsec were thus far examined in numerous pre-clinical setups of experimental tissue damage, including acute myocardial infarction, autoimmune myocarditis, stroke, spinal cord injury amongst others (for review see [19]). Topical administration of PBMCsec enhanced wound healing in a murine full-thickness skin model [16] and improved tissue survival in a rodent epigastric flap model [20]. Similar results were observed in a porcine model of burn injury, where the application of PBMCsec markedly improved epidermal thickness after injury [21]. In addition, beneficial effects of PBMCsec were also observed after intraperitoneal or intravenous administration. PBMCsec decreased the affected area and improved neurological outcome in rodent models of cerebral ischemia and acute spinal cord injury after systemic application [22,23].

Furthermore, PBMCsec ameliorated myocardial damage, improved overall cardiac performance and promoted cytoprotection of cardiomyocytes in rodent and porcine models of acute myocardial infarction (AMI) [10,14]. Interestingly, transcriptional changes were not restricted to the infarcted myocardium and the circumjacent heart tissue but also detected in the liver and the spleen, suggesting a systemic effect of PBMCsec beyond the site of injury [24].

For the treatment of cardiovascular pathologies caused by thromboembolic occlusion of arterial blood flow such as acute myocardial infarction and ischemic stroke, the rapid interventional re-establishment of vessel perfusion still remains the first-line therapy as it maximizes rescue of ischemic tissue and thus decisively determines primary outcome in affected patients [25]. However, increasing emphasis is attributed to the attenuation of damages secondary to reperfusion injury, as it represents a considerable risk factor for long-term tissue functions [26]. During extended periods of oxygen deprivation, cells suffer from intracellular calcium overload [27] and disturbed mitochondrial energy production [28], which ultimately lead to cell death and the release of inflammatory mediators and reactive oxygen species [29]. In turn, inflammatory cells infiltrate the damaged area and release pro-inflammatory cyto- and chemokines. In addition, neutrophils release serine-proteases [30,31], that contribute to endothelial barrier dysfunction, further enhancing vascular injury in small arterial blood vessels and downstream capillaries [32]. As a result, large amounts of intravascular fluids along with the damaging mediators diffuse into the interstitial space to cause further damage [32].

Since PBMCsec showed tissue regenerative properties in several animal models and there is an urgent need for novel systemic tissue-regenerative therapeutic interventions, we investigated the effects of PBMCsec on white blood cells and on the endothelial barrier function.

## Results

### PBMCsec modulates the gene signature of T cells, B cells, granulocytes, and monocytes

To investigate the degree to which PBMCsec alters the transcriptional landscape of immune cells in human whole blood, we conducted single cell RNA sequencing (scRNAseq) of whole blood samples treated *ex vivo* with PBMCsec for 8 hours. A methodological overview of the experimental approach employed in this study is provided in Fig.1. Bioinformatics analysis and UMAP-clustering revealed four main cell populations consisting of monocytes, T-cells, B-cells and granulocytes in all investigated conditions (Fig. 2A). Identification was based on expression of cluster marker genes including CD14 Molecule (*CD14)*, CD3d Molecule (*CD3D)*, Membrane Spanning 4-Domains A1 (*MS4A1)* and Peptidase Inhibitor 3 (*PI3)* for the respective cell types (Fig. S1). Treatment with PBMCsec did not result in a significant change in relative cell numbers (percentage of cells in untreated vs PBMCsec for T-cells: 75.73% vs 74.86%; monocytes: 14.31% vs 14.97%; B-cells: 8.89% vs. 8.09%; granulocytes 1.03% vs 1.90%) (Fig. 2B). The transcriptional heterogeneity between celltypes was confirmed in a heatmap showing the average expression of cluster-defining genes of each cell type (Fig.2C). Next, we assessed changes in gene expression for all cell populations after treatment with PBMCsec. We calculated the number and distribution of significantly up-(Fig. 2D) and downregulated genes (Fig. 2E) for each cell-type compared to the untreated control. A total number of 45 differentially expressed genes (DEG) (16 upregulated; 29 downregulated) were detected in T-cells, 72 in B-cells (28 upregulated, 44 downregulated) and 17 in granulocytes (2 upregulated; 15 downregulated) (SFig. 2A-C). Monocytes displayed the highest number of differentially expresses genes (173 upregulated; 148 downregulated), including upregulation of Serpin Family B Member 2 (*SERPINB2*), Epiregulin (*EREG*), Interleukin 1 Beta (*IL1B*), C-X-C Motif Chemokine Ligand 1 (*CXCL1*), C-X-C Motif Chemokine Ligand 3 (*CXCL3)*, (*CXCL5)*, and Vascular Endothelial Growth Factor A (*VEGFA*), and downregulation of S100 Calcium Binding Protein A8 (*S100A8*), S100 Calcium Binding Protein A9 (*S100A9*), Allograft Inflammatory Factor 1 (*AIF1*), CD36 Molecule (*CD36*) and CD163 Molecule (*CD163*) amongst others (Fig. 2F). From this we conclude that *ex vivo* stimulation of human whole blood with PBMCsec significantly changes gene expression in blood immune cells, most prominently in monocytes.

**Figure 1.**
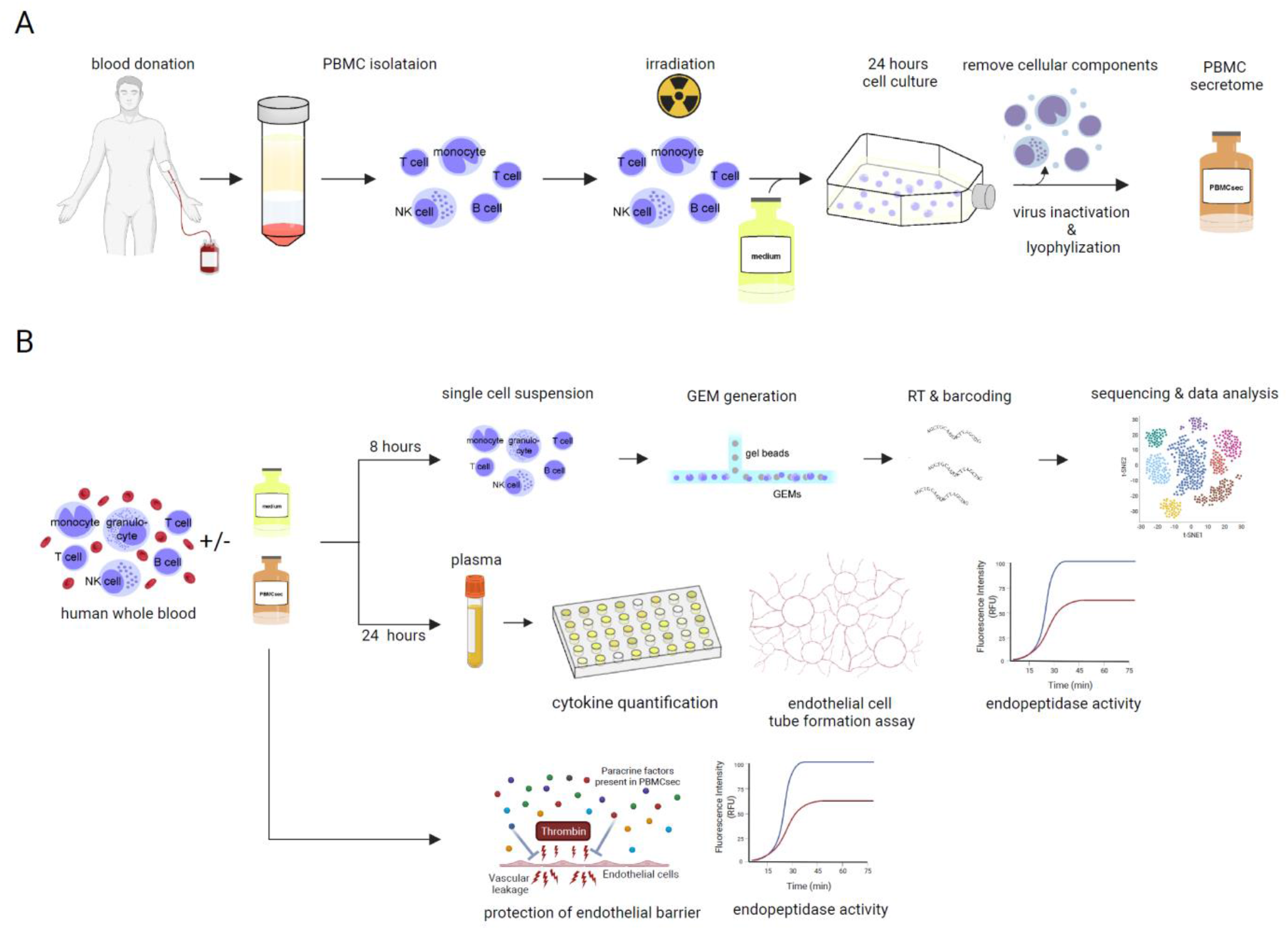
Experimental overview. (A) Isolation of PBMCs from healthy donors and subsequent procedures necessary for the generation of PBMCsec. (B) The experimental setup involved two branches. In order to investigate pharmacodynamics effects of PBMCsec human whole blood cells were treated with PBMCsec or left untreated for 8 hours prior to further processing them for single cell RNA sequencing. Plasma was collected from PBMCsec treated whole blood and controls after 24-hour long incubation and analyzed in a series of functional assays. In addition, we evaluated effects mediated directly by PBMCsec on endopeptidase activity and endothelial barrier protection. This figure was illustrated using Biorender.com.

**Figure 2.**
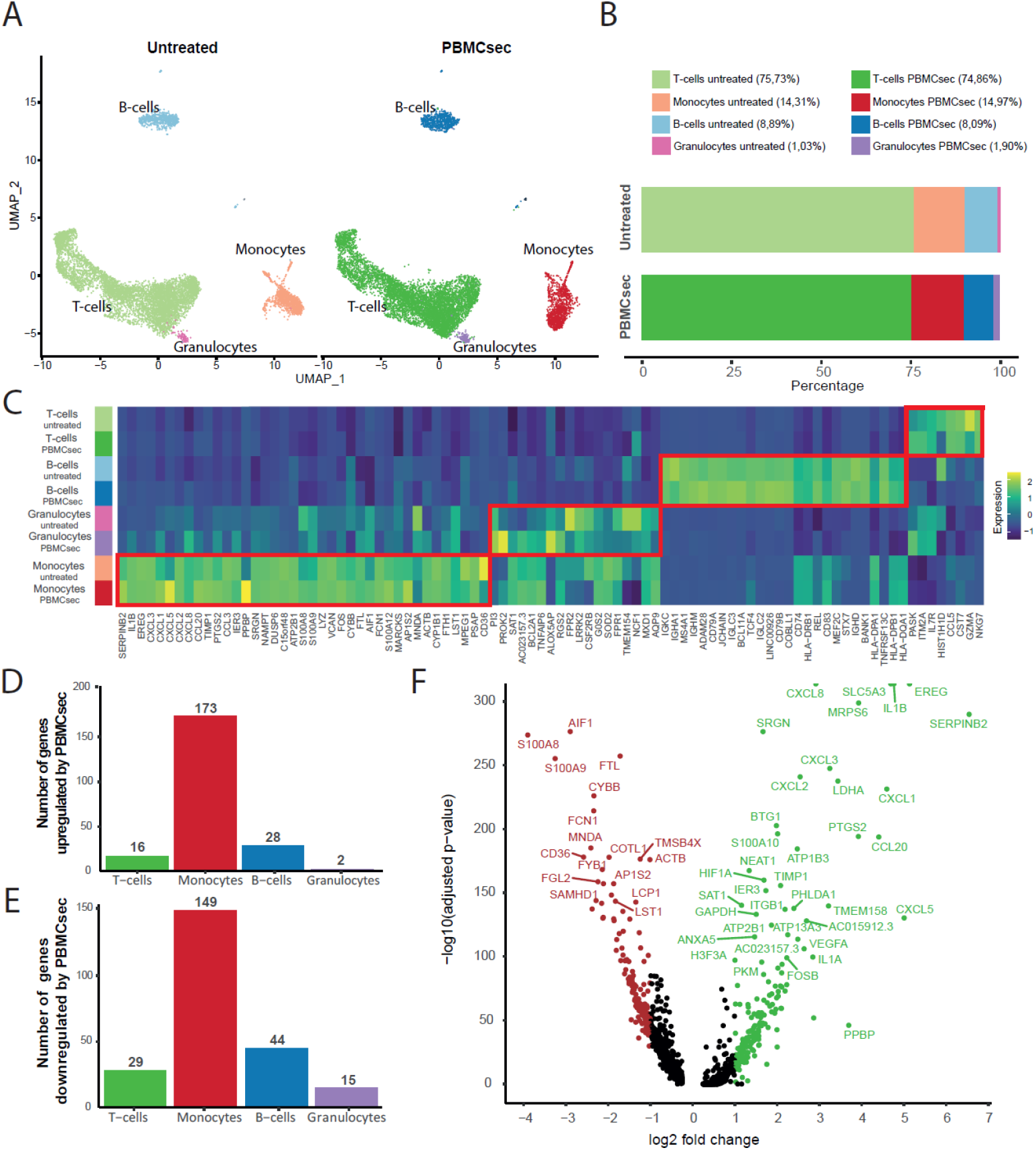
Ex vivo treatment of human whole blood cells alters the transcriptional profile in the Monocyte subset and upregulates pathways associated with tissue-regeneration. (A) UMAP-clustering identifies T-cells, monocytes, B-cells and granulocytes in untreated and PBMCsec-treated samples with (B) similar cell frequency for each cluster between the investigated conditions. (C) Heat-map of cluster-defining marker genes of normalized gene expression shows distinct gene patterns between T-cells, Monocytes, B-cells and Granulocytes. Barplots represent the number of (D) up - and (E) downregulated genes in PBMCsec-treated samples compared to untreated control. Genes with an adjusted p-value < 0.05 and an average Log_2_ foldchange ≥ 1 or ≤-1 were considered as DEG. (F) Volcano plot showing the regulated genes in monocytes treated with PBMCsec when compared to untreated monocytes. Upregulated genes are marked in green, while downregulated genes are shown in red.

### PBMCsec induces tissue-regenerative pathways in monocytes from human whole blood

Next, we identified biological pathways associated with the differentially regulated genes across the identified cell types. Biological functions such as angiogenesis and cytokine production were enriched in T- and B cells treated with PBMCsec (Fig. S2D and Fig. S2F) while activation of myeloid cells and responses to oxidative stress were associated with the downregulated gene sets (Fig. S2E and Fig. S2G). No significantly regulated pathways were identified in granulocytes. The top Gene Ontology pathways associated with upregulated genes in monocytes after treatment with PBMCsec included response to lipid and to interleukine-1 along with terms strongly associated with wound healing, angiogenesis, regulation and production of cytokines and regulation of endopeptidase activity (Fig. 3A). Major pathways associated with biological processes resulting from the downregulated gene set in monocytes treated with PBMCsec involved activation and differentiation of leukocytes, generation of reactive oxide species and processing and presentation of antigens amongst others (Fig. 3B). The complete lists of all up- and downregulated pathways (Fig. S3A and Fig. S3B) and genes (supplementary file 1 and supplementary file 2, respectively) are provided as supplementary information. A closer look at the genes involved in the degranulation and cell activation of leukocytes revealed different members of the S100 and leukocyte immunoglobulin-like receptor (LILR) families to be downregulated in monocytes treated with PBMCsec (Fig. S4A). Furthermore, we observed a downregulation of genes associated with the generation of reactive oxygen species including the scavenger receptor CD36, the pattern recognition receptor CLEC7A, a subunit of the NADPH oxidase complex in NCF1 and others (Fig S4B). We further sought to confirm our findings using ClueGO to investigate molecular functions related to these gene sets. In line with the initial analysis, we again identified strong associations with molecular functions related to cyto- and chemokine activity, signaling and negative regulation of cysteine-type endopeptidase activity for the upregulated genes (Fig. 3C), while functions related to immune receptor signaling, oxidoreductase activity and antigen presentation were significantly associated with the set of downregulated genes (Fig. 3D). From this analysis, we conclude that monocytes are mostly affected by PBMCsec, resulting in the induction of tissue-regenerative processes while processes associated with leukocyte activation and reactive oxygen species generation are downregulated.

**Figure 3.**
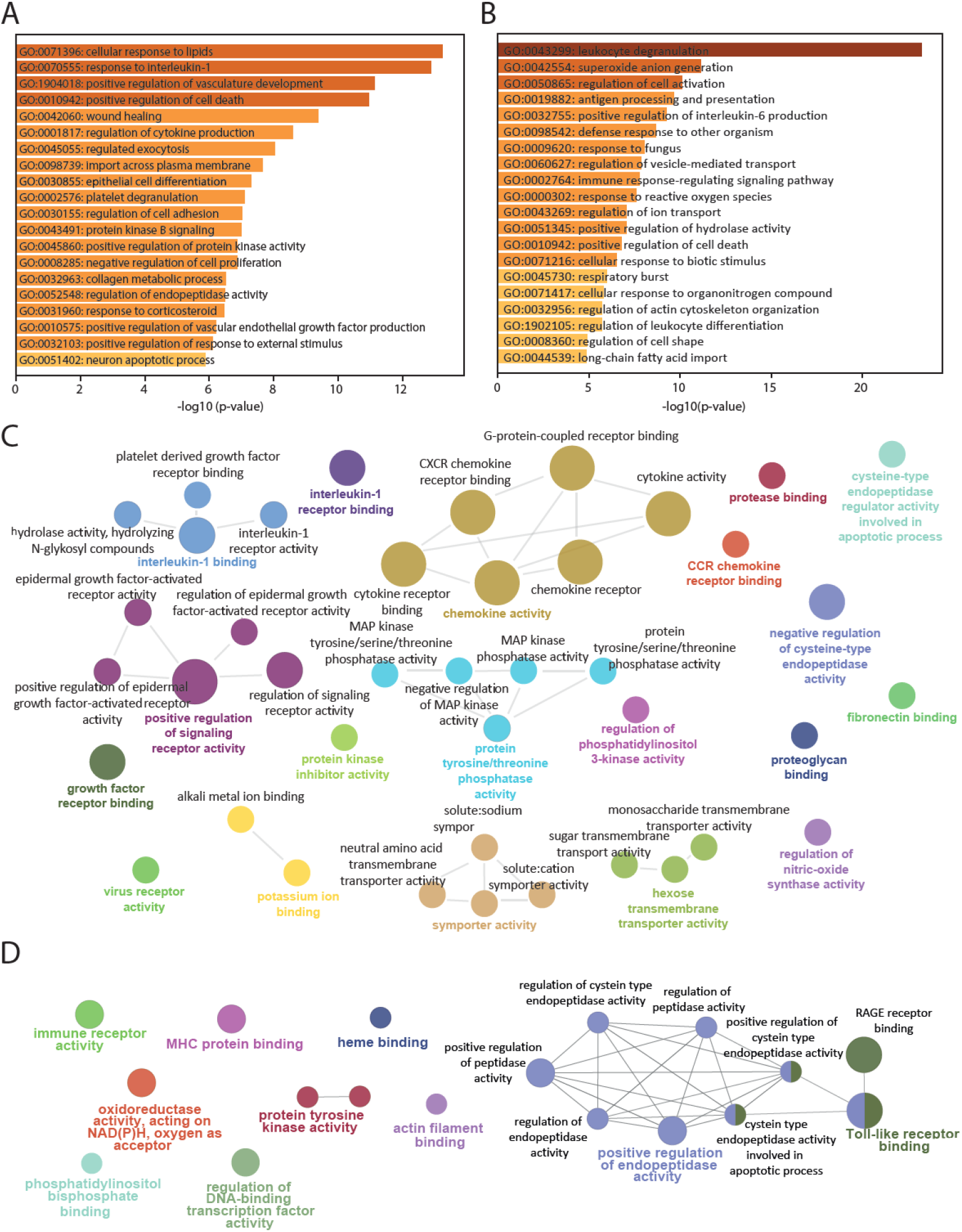
Tissue-regenerative pathways are upregulated in monocytes treated with PBMCsec. Gene ontology pathway analysis was done in Metascape. The top 20 biological processes affected by the upregulated (A) and downregulated (B) DEG in monocytes treated with PBMCsec are listed and hierarchically sorted by –log_10_(p-value) and enrichment factor (>2). (C) ClueGO visualization of molecular functions associated with upregulated and (D) downregulated genes in monocytes treated with PBMCsec.

### Paracrine factors in the plasma of PBMCsec-treated whole blood exert pro-angiogenetic effects *in vitro*

Since several cytokines and growth factors were upregulated in monocytes treated with PBMCsec (Fig. 2F), we next assessed the protein levels of selected factors in plasma derived from PBMCsec-treated whole blood. PBMCsec, fresh plasma collected immediately after venipuncture, and untreated plasma after *ex vivo* incubation for 24 hours served as controls. Gene expression of *CXCL1, CXCL5* and *VEGFA* showed strong upregulation in monocytes after stimulation with PBMCsec (Fig. 4A). This was confirmed on protein level, when CXCL1 (2873 ± 2093 pg/ml), CXCL5 (6413 ± 1911 pg/ml) and Vegf-A (117.2 ± 13.04 pg/ml) were strongly elevated in plasma PBMCsec, while being undetectable in controls (Fig. 4B). In contrast, several pro-inflammatory factors, such as IL-1β, TNF-α and IFN-γ were not detectable in the plasma of PBMCsec treated whole blood (data not shown). We next aimed to assess the pro-angiogenic capacity of fresh plasma, plasma of untreated and plasma of PBMCsec-treated whole blood in an *in vitro* endothelial cell tube formation assay (Fig 4C). While controls showed little ability to induce tube formation, plasma of PBMCsec-treated whole blood displayed the highest pro-angiogenic properties as indicated by a significant increase in total segment length as well as in numbers of nodes and junctions (Fig. 4D). In summary, we could demonstrate that expression and secretion of pro-angiogenic paracrine factors is strongly increased in whole blood cells following treatment with PBMCsec.

**Figure 4.**
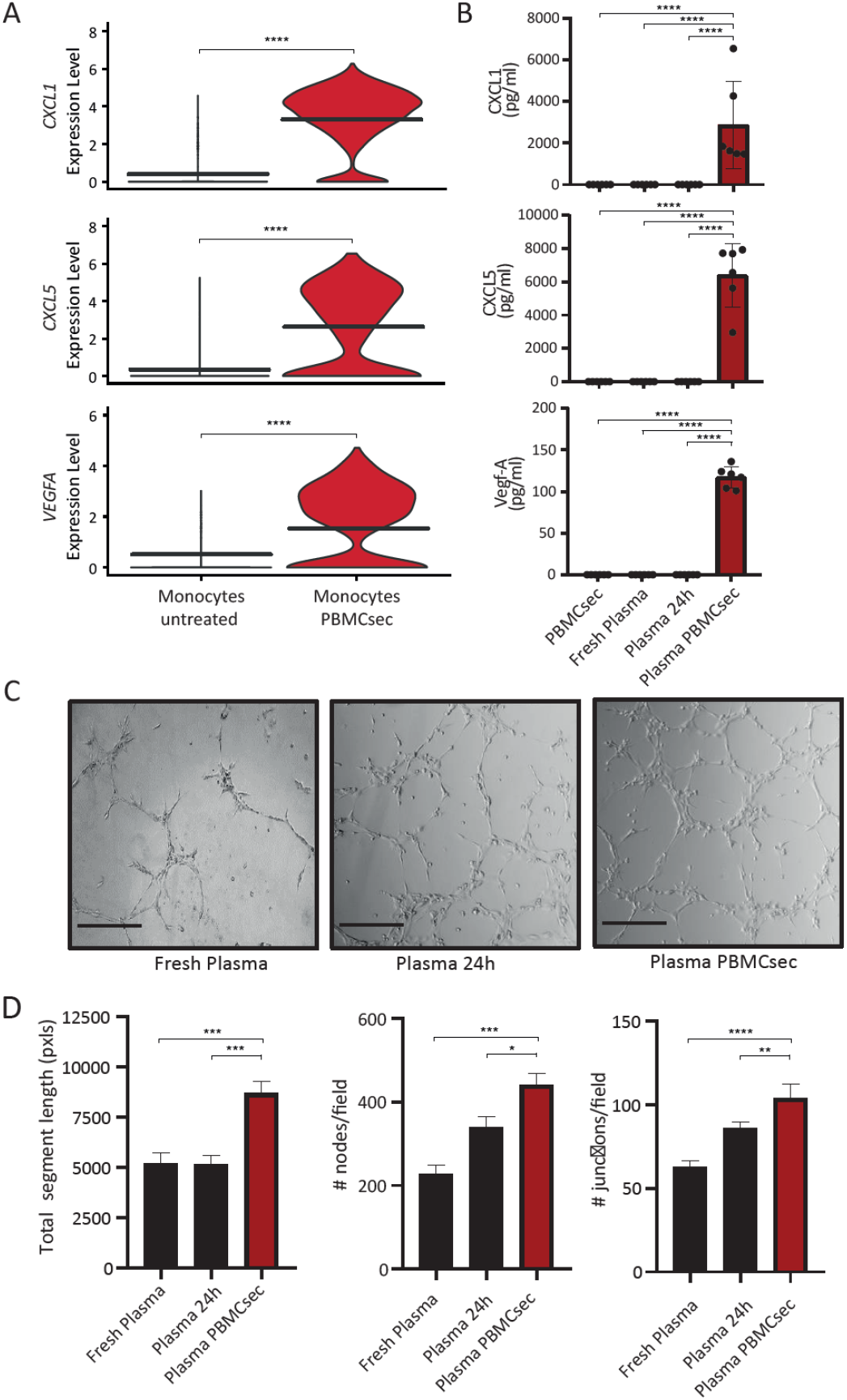
Paracrine factors are induced by PBMCsec in human plasma and enhance endothelial cell tube formation in vitro. (A) Gene expression of the pro-angiogenetic chemokines *CXCL1* and *CXCL5* as well as growth factor *VEGFA* are strongly induced in monocytes treated with PBMCsec when compared to untreated monocytes (B) Assessment of protein levels for CXCL1, CXCL5 and Vegf-A by ELISA in plasma from PBMCsec-treated whole blood and controls (n = 6). (C) Representative images of HUVECs tube formation assay in presence of fresh plasma, plasma from untreated whole blood and plasma obtained from PBMCsec-treated whole blood after 24-hour *ex vivo* cultivation (scale bar 250μm) with (D) analysis of number of nodes and junctions per field and total segment length (n=3). Ordinary one-way ANOVA was performed. Dunnett’s multiple comparison test was carried out to compare groups. * indicates p-value < 0.05; ** indicate p-value < 0.01; *** indicate p-value < 0.001; **** indicate p-value < 0.0001

### PBMCsec inhibits protease activity *in vitro* and induces the selective serine-protease SERPINB2 in human whole blood *ex vivo*

As our pathway analysis also revealed an association of upregulated genes in monocytes with the regulation of endopeptidase activity, we next aimed to confirm this finding in functional assays *in vitro*. Therefore, we performed a protease activity assay to test a potential anti-proteolytic effect of PBMCsec on the non-selective serine-protease trypsin. While medium control showed a negligible inhibitory effect on protease activity (3.72 ± 0.55 %), PBMCsec lead to a significant inhibition of the enzymatic activity (40.51 ± 4.728 %) (Fig. 5A). Furthermore, *SERPINB2* displayed the highest positive log_2_fold-change induction (Fig 5B) of all DEGs in monocytes treated with PBMCsec in our sequencing analysis. *SERPINB2* is a member of the serpin superfamily of serine proteases and encodes plasminogen activator inhibitor type II (PAI-2) which is involved in the irreversible inhibition of urokinase [33]. First, we determined the protein levels of SERPINB2 in PBMCsec, fresh plasma, plasma of untreated whole blood and compared them to plasma of PBMCsec-treated whole blood. In accordance with the present literature, only very low levels of SERPINB2 were detectable in the control samples. In contrast, plasma obtained from PBMCsec-treated whole blood showed significantly elevated levels of SERPINB2 (11908 ± 3530 pg/ml) (Fig. 5C). As SERPINB2 mainly acts as an inhibitor for urokinase [33], we next evaluated plasma levels of urokinase as well as the inhibitory effect of plasma PBMCsec on its activity *in vitro*. Whereas high levels of urokinase were still detected (Fig 5D), urokinase activity was strongly reduced by PBMCsec-treated plasma (Fig. 5E), indicating that the observed urokinase inhibitory action was a result of the presence of a urokinase inhibitor rather than a quantitative decrease in available urokinase. While *in vitro* urokinase activity was not affected by PBMCsec alone, addition of fresh plasma (38.45 ± 5.94%) and plasma of untreated whole blood (40.78% ± 17.07%) resulted in a considerable inhibition of urokinase activity. Interestingly, this effect was even more pronounced in the presence of plasma obtained from PBMCsec-treated whole blood (79.48 ± 2.15%) (Fig 5E). Together, these data show that soluble factors with anti-proteolytic activities are present in PBMCsec or strongly induced in white blood cells after treatment with PBMCsec.

**Figure 5.**
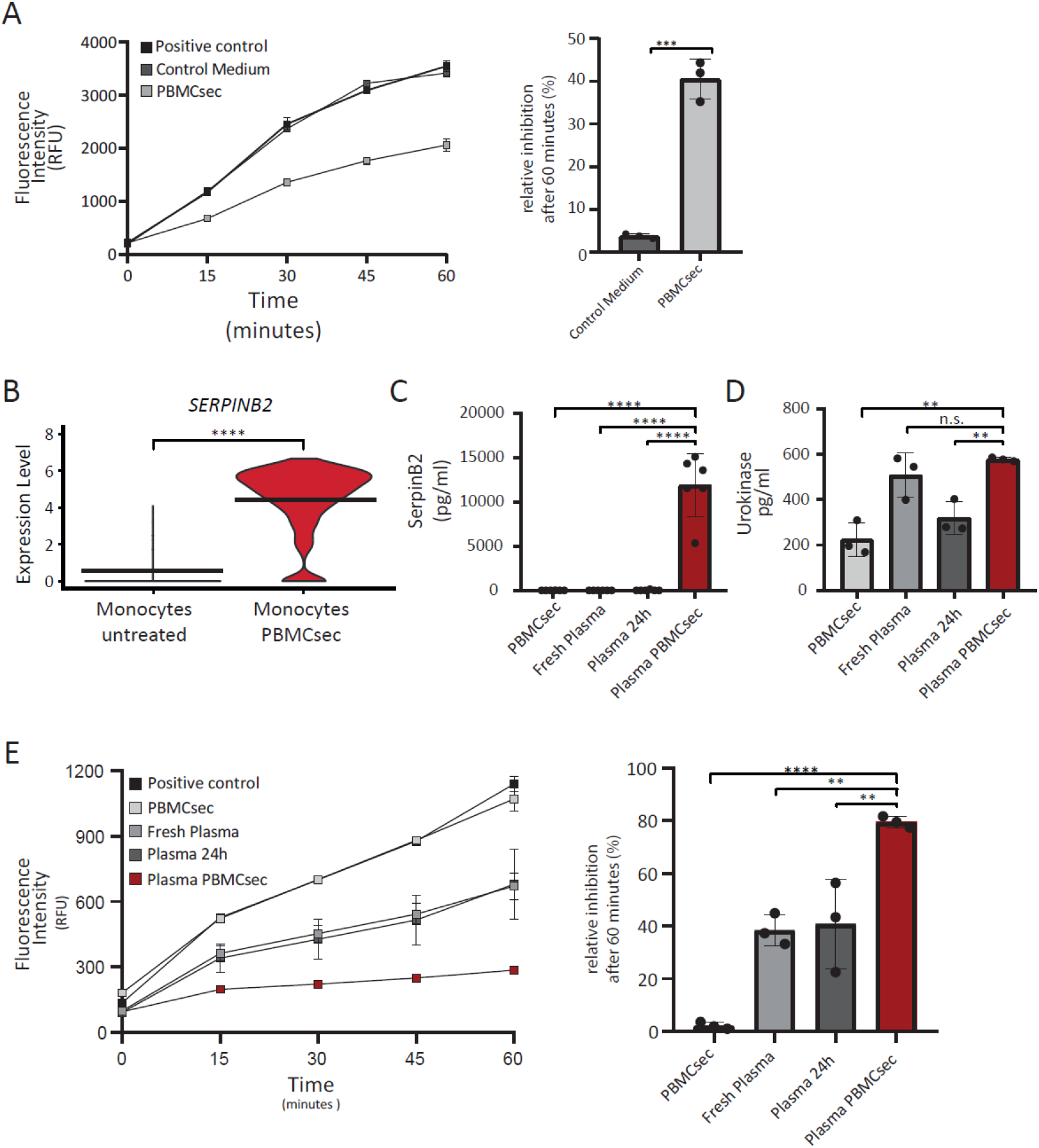
PBMCsec inhibits serine-protease activity in vitro and increases levels of SERPINB2 in plasma of PBMCsec-treated whole blood. (A) Trypsin activity measured by increase in fluorescence intensity of cleaved substrate over time in the presence and absence of PBMCsec (n=3). (B) Gene expression of the urokinase - inhibitor *SERPINB2* in monocytes treated with PBMCsec (p-value: <0.0001) compared to untreated monocytes. (C) SerpinB2 and (C) urokinase protein levels of fresh plasma, untreated plasma and plasma of PBMCsec-treated whole blood. (E) In vitro assessment of urokinase activity of fresh plasma, plasma from untreated whole blood and plasma obtained from PBMCsec-treated whole blood. * indicates p-value < 0.05; ** p-value < 0.01; *** indicate p-value < 0.001; **** indicate p-value < 0.0001

### PBMCsec ameliorates thrombin-induced decrease in endothelial barrier function

On the one hand, the activity of inflammatory mediators and serine-proteases supports angiogenesis, on the other hand it may also increase vascular leakage by decreasing the endothelial barrier function [34,35]. Hence, we examined whether PBMCsec affected thrombin-induced drop in endothelial barrier function by utilizing electrical cell-substrate impedance sensing. Addition of thrombin resulted in a transient decrease of trans-endothelial resistance, reaching the lowest point 5 minutes after stimulation (29 ± 12% barrier function relative to untreated basal medium control) (Fig. 6A). While the simultaneous addition of thrombin and control medium resulted in a similar change in barrier resistance (38 ± 2%) compared to basal medium, PBMCsec inhibited thrombin-induced decrease in endothelial resistance (84 ± 14%) (Fig. 6A). Therefore, PBMCsec positively influences endothelial barrier function in vitro by attenuating thrombin-mediated changes in endothelial cells.

**Figure 6.**
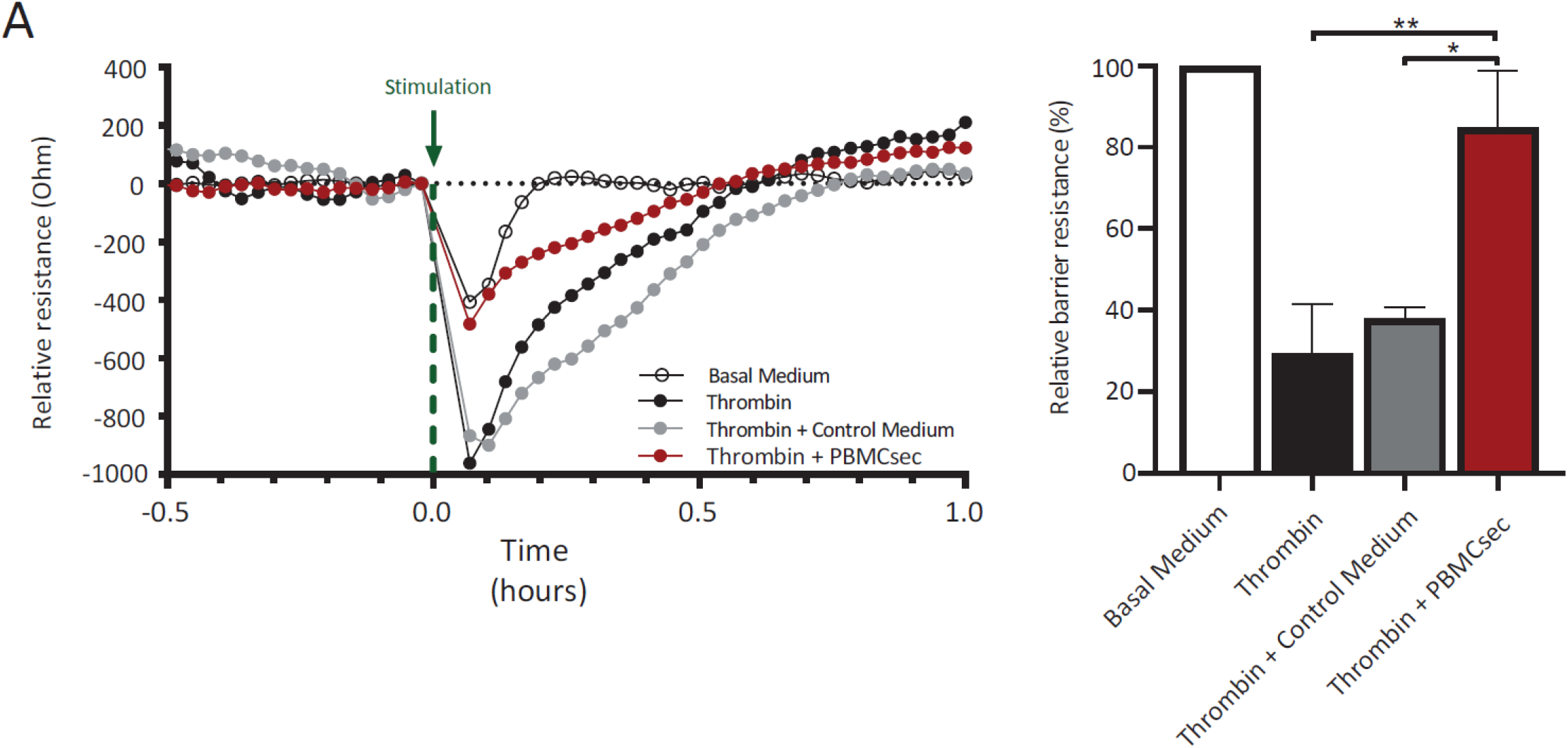
PBMCsec amerliorated thrombin-induced decrease in endothelial cell barrier function. (A) Representative evaluation of one of three independently conducted experiments. Paracellular resistance at 250Hz was measured in DMECs challenged with 2 units/ml Thrombin alone or in combination with control medium or PBMCsec at 12.5 units/ml over time. The green arrow marks the time point of stimulation with the respective treatments. Barplot shows the changes in endothelial barrier function for the investigated conditions relative to the basal medium control. One-way ANOVA was performed with post -hoc Dunnett’s multiple comparison test to compare groups to basal medium. *indicates p-value < 0.05; ** indicate p-value < 0.01

## Discussion

The beneficial effects of PBMC-derived cell secretomes in the regeneration of damaged tissues and organs have already been well described [10,14,17,19,22,23,36]. In order to transit from these promising preclinical studies to the treatment of patients, the safety and tolerability of topical administration of autologous PBMCsec in dermal wounds was successfully assessed in a clinical phase I trial (MARSYAS I, NCT02284360) [37]. Based on these results, an ongoing clinical phase I/II trial has been initiated to evaluate the safety and efficacy of topically administered allogeneic PBMCsec for the treatment of diabetic foot ulcers (MARSYAS II, NCT04277598) [38]. In the light of future therapeutic approaches using systemic administration of PBMCsec to regenerate inner organs, we analyzed in the present study the effects of PBMCsec on whole blood and the endothelium in more detail.

Using scRNAseq, we identify a strong induction of tissue-regenerative pathways after treatment of human whole blood with PBMCsec *ex vivo*. Our analysis revealed that changes of the transcriptional profile of white blood cells treated with PBMCsec were most pronounced in monocytes, while all other celltypes showed comparably low alterations in gene expression. Our finding is in line with previous publications where functional changes were mainly reported in monocytes after stimulation with conditioned media from MSCs [39][40]. As monocytes are an important part of the innate immune system, a rapid response to extracellular stimuli, such as PBMCsec, was expectable. Interestingly, only few granulocytes, which also represent a main constituent of innate immunity, were detected in our single cell analysis. A low detectability of the different granulocyte populations in scRNAseq has been reported before and might be due to their overall low RNA content and presence of RNAses [41]. This assumption is further supported by the extremely low mRNA counts in the few granulocytes detected in our analysis. Interestingly, also T and B-cells showed only minor gene regulation after exposure to the secretome. As both cell types belong to the adaptive branch of the immune system, it is tempting to speculate that repeated exposure of lymphoid cellular subsets to PBMCsec might result in a more pronounced adaptive immune response. Further studies will be necessary to fully address this question.

Examination of the altered gene sets in monocytes revealed upregulation of several members of the chemokine (C-X-C motif) ligand (CXCL) family (CXCL1, CXCL2, CXCL3, CXCL5 and CXCL8), as well as different growth factors after stimulation with PBMCsec. In addition, high protein levels of CXCL1, CXCL5 and VEGF-A were detected in the plasma of PBMCsec-treated whole blood. These factors have previously been described as important constituents of PBMCsec, contributing to its tissue regenerative actions [14,42]. Yet, we were not able to detect these proteins in PBMCsec in our assays. This discrepancy could be explained by the lower concentration of PBMCsec that we used in our study. Nevertheless, all these factors were strongly induced in whole blood and abundantly present in the plasma derived from PBMCsec-treated whole blood, indicating a significant amplification of the production of tissue regenerative factors. In addition, our transcriptional analysis revealed a downregulation of genes involved in the generation of superoxide anion species with an inhibitory effect on the formation of reactive oxygen species in PBMCsec-treated monocytes. Expression of the scavenger receptor CD36 on monocytes treated with PBMCsec was amongst the most strongly down-regulated genes. In monocytes and macrophages, activation of this receptor promotes increased production of reactive oxygen species which in turn reinforce damages to the vasculature [43]. Furthermore, CD36 is known to modulate the uptake of oxidized lipid species by macrophages, leading to functional changes in these cells. As they accumulate lipids, these then so-called foam cells promote proliferation and inflammation of endothelial and smooth muscle cells, which contributes to the formation of atherosclerotic plaques [44]. Given that reactive oxygen species increase after re-establishment of perfusion and aggravate tissue damage in a series of cardiovascular pathologies, our data offer a potential mechanism by which PBMCsec might directly and/or indirectly reduce tissue damage after ischemic conditions. Future studies will elucidate the influence of PBMCsec on the generation and amelioration of reactive oxygen species.

Previous work from our group has demonstrated that paracrine factors released from γ-irradiated PBMCs positively influence tissue regeneration by affecting endothelial cell survival and sprouting [11]. In addition, we here showed that PBMCsec was also able to inhibit thrombin-induced damage of the endothelial barrier. This effect might counteract increased leakage of blood vessels, thereby preventing insufficient perfusion of the vital part of the injured organ and amplification of the damage. We also observed an inhibitory effect of the plasma of PBMCsec-treated whole blood on the serine-protease urokinase, which was not detected in PBMCsec alone. Thus, *de novo* synthesis of the urokinase inhibitor SERPINB2 could mediate this effect. Serine-protease inhibitors are widely discussed as promising therapeutic approaches for several disease entities. Recently, Vorstandlechner et al. identified dipeptidyl-peptidase 4 (DPP4) and urokinase (PLAU) as important drivers of skin fibrosis and targeted inhibition of both molecules was shown to reduce myo-fibroblast formation and improve scar quality [45]. Implications for therapeutic targeting of serine-proteases are also evident for myocardial infarction [46,47]. While early re-vascularization of occluded blood vessels still remains the gold standard intervention to maximize rescue of functional tissue, increasing emphasis is put on the reduction of late onset events caused by accumulation of pro-inflammatory stimuli, proteases and other detrimental factors, secondary to reperfusion injury [30]. Mauro and colleagues reported reduced infarction sizes following treatment with Alpha-1-antitrypsin in addition to revascularization in a murine model of myocardial infarction [48]. This was further reinforced by a report by Hooshdaran et al. where inhibition of the serine proteases cathepsin G and chymase were shown to reduce adverse cardiac remodeling, myocyte apoptosis and fibrosis in a murine model of myocardial ischemia reperfusion injury [49]. In addition, recent publication by Sen and colleagues reported a central role for SerpinB2 in the coordinated resolution and repair of damaged tissue after ischemia-reperfusion in a murine kidney injury model [50]. These data suggest that in addition to the protective action of serine-protease inhibitors investigated in our study, they might also show valuable tissue-regenerative and anti-fibrotic properties. Further studies are needed to fully decipher the contribution of protease inhibitors present in PBMCsec, to the inhibition of tissue-damage or the restoration of damaged tissue and organs.

Paracrine signaling for the prevention of a pro-inflammatory response and cell death as well as induction of angiogenesis are essential to restore damaged tissues and organs [40][51]. We have previously shown that PBMCsec was able to positively modulate all of these events [11,14,52] and administration of a single dose of PBMCsec was sufficient to almost completely inhibit heart damage in a porcine model of experimental myocardial infarction [14]. However, given that pharmacodynamics investigations revealed that several main components of PBMCsec were only traceable for minutes to a maximum of 5 hours in the blood of rats and dogs after intravenous application of human PBMCsec, this finding was rather unexpected [53]. Whether human PBMCsec induced a comparable release of pro-regenerative factors in these animal models has not been investigated so far. Our new data provide a reasonable explanation for this observation and suggest that PBMCsec induces the production of a new secretome in human white blood cells with additional regenerative properties. As a result, the pharmacodynamics effect of a single application of PBMCsec can be significantly prolonged by continuous stimulation, comparable to the behavior of a damped wave. This secretome-induced secretome, produced over an extended period of time, might therefore multiply the pro-regenerative properties of PBMCsec by combining them with those of the newly induced factors. However, further studies in a human setting will be necessary to fully explore the tissue-regenerative potential of the newly formed secretome and to examine for how long this effect can be maintained.

In summary, our scRNA-seq analysis identified key mechanisms, potentially contributing to tissue regeneration, which might occur after systemic application of a cell secretome derived from irradiated PBMCs. PBMCsec displays a broad spectrum of mechanistic modes of actions that greatly complement each other and therefore positively contribute to tissue regeneration in a wide range of pathological settings. In addition, our data suggest that the effects *in vivo* are not restricted to direct actions of PBMCsec but also arise from further stimulation of circulating monocytes in the blood. Further human clinical studies in the future will clarify the underlying mechanisms and the therapeutic benefit of systemic treatment of damaged organs with PBMC-derived secretomes.

## Materials and Methods

### Ethics statement

This study was conducted in accordance with the Declaration of Helsinki and local regulations. Blood samples were obtained from healthy volunteers who had given their consent to donate. Use of primary HUVECs, primary DMECs and blood samples was approved by the Institutional Review Board of the Medical University of Vienna (Ethics committee votes: 1280/2015, 1539/2017 and 1621/2020). All donors provided written informed consent.

### Generation of PBMCsec

Isolation of PBMCs and generation of PBMCsec have previously been described in detail [54] and a graphical overview is given in Fig. 1A. In brief, PBMCs were enriched by Ficoll-Paque PLUS (GE Healthcare, Chicago, IL, USA) density centrifugation, diluted to a concentration of 2.5 × 10^7^ cells/mL in CellGenix granulocyte-monocyte progenitor dendritic cell medium (CellGenix, Freiburg, Germany) and exposed to 60 Gy Caesium 137 γ-irradiation (IBL 437C, Isotopen Diagnostik CIS GmbH, Dreieich, Germany). Following 24-hour long incubation, cells and cellular debris were removed by centrifugation with 800g for 15 minutes and supernatants were passed through a 0.2 mm filter. The cell-free secretome generated by 2.5 × 10^7^ cells/mL corresponds to 25 units/ml PBMCsec. Next, methylene blue treatment was performed for viral clearance [44]. Secretomes were lyophilized, terminally sterilized by high-dosage-irradiation (Gammatron 1500, UKEM60Co irradiator with a maximum capacity of 1.5 MCi), and cryopreserved. Lyophilized compounds were reconstituted in 0.9% NaCl (B. Braun Melsungen AG, Melsungen, Germany) to the desired concentrations.

### Preparation of single cell suspension of human whole blood

For scRNAseq, heparinized human whole blood was drawn from two age-matched male donors. A total of three ml of whole blood was either treated with PBMCsec (GMP APOSEC lot number: A00918399135; diluted in 0.9% NaCl; final concentration: 12.5 units/ml) or left untreated. Samples were incubated at 37°C for 8h. Red blood cells were removed by Red Blood Cell Lysis Buffer (Abcam, Cambridge, USA). Cells were then washed twice with PBS containing 0.04% bovine serum albumin (BSA) and sequentially passed through 100- and 40-μm cell strainer. Using the LUNA-FL Dual Fluorescence Cell Counter (BioCat, Heidelberg, Germany) and the Acridine Orange/Propidium Iodide Cell Viability Kit (Logos Biosystems, Gyeonggi-do, South Korea) samples were set at a concentration of 1×10^6^ cells/ml and displayed a viability above 90%.

### Gel Bead-in Emulsion (GEMs) - and library preparation

Single-cell RNA-seq was performed using the 10X Genomics Chromium Single Cell Controller (10X Genomics, Pleasanton, CA, USA) with the Chromium Single Cell 3′ V3 Kit following manufacturer’s instructions. After quality control, RNA sequencing was performed by the Biomedical Sequencing Core Facility of the Center for Molecular Medicine (Center for Molecular Medicine, Vienna, Austria) on an Illumina HiSeq 3000/4000 (Illumina, San Diego, CA, USA). For donor 1, we detected 2003 cells in the untreated sample and 1281 cell in the PBMCsec-treated sample, while donor 2 had 12356 cells in the untreated sample and 10865 in the PBMCsec treated sample. Raw sequencing data were then processed with the Cell Ranger v3.0.2 software (10X Genomics, Pleasanton, CA, USA) for demultiplexing and alignment to a reference genome (GRCh38).

### Data analysis

Secondary data analysis was performed using R Studio in R (Version 4.0.4; The R Foundation, Vienna, Austria) using the R software package “Seurat” (Seurat v.4.0.0, Satija Lab, New York, NY, USA). Cells were first analyzed for their unique molecular identifiers (UMI) and mitochondrial gene counts to remove unwanted variations in the scRNAseq data. Cells with UMI counts below 200 or above 2500 and more than 10% of mitochondrial genes were excluded from the data set. Next, we followed the recommended standard workflow for integration of scRNAseq datasets [55]. Data were scaled and principal component analysis (PCA) was performed. Statistically significant principal components (PCs) were identified by visual inspection. Using the Louvain algorithm at a resolution of 0.025, we identified a total of 4 communities. The preselected PCs and identified clusters served for Uniform Manifold Approximation and Projection for Dimension Reduction (UMAP). After bioinformatics integration of datasets of untreated and PBMCsec-treated samples, erythrocytes were removed by excluding all cells with expression of Hemoglobin subunit beta (HBB) > 0.5. Clusters were then annotated based on the expression of well - established cell-type-defining marker genes. We used UMAP-plots, feature plots, heat maps, volcano plots and violin plots to visualize differences between the investigated conditions. To determine DEGs, normalized count numbers were used. We applied the FindMarkers argument using default settings to calculate DEGs for clusters of interest between conditions with a log-foldchange threshold of 0.25 and an adjusted p-value < 0.05. A Log2-fold-change increase of gene expression above 1 was considered as upregulation while a decrease below -1 was considered as downregulation. Only genes with an avgLog2FC above 1 and below -1 were forwarded to the Metascape [56] online software package to identify significantly enriched pathways (-log10(p-value) > 2). Additionally, the same gene sets were processed by Cytoscape plug-in ClueGO to visualize significantly (p-value < 0.05, kappa score: 0.4) enriched molecular functions for the investigated conditions.

### Protein quantification by Enzyme-linked Immunosorbent Assay (ELISA)

For *in vitro* experiments, heparinized human whole blood was drawn from male donors. A total of 3 ml of whole blood was centrifuged (1000g for 10 minutes at room temperature) freshly after venipuncture or after 24-hour long cultivation of whole blood in absence or presence of 12.5 units/mL PBMCsec. Plasma samples were then stored at -20°C until further use. Protein levels of Human CXCL1, human CXCL5, human SERPINB2, human VEGF-Aand human Urokinase (R&D Systems, Biotechne, Minneapolis, USA) were quantified by ELISA as recommended by the manufacturer. Absorbance was measured at 450 nm by a Spark multimode microplate reader (Tecan, Männedorf, Switzerland) and analyte quantifications were determined using external standard curves.

### Protease activity assays

To test the inhibitory effects of PBMCsec on protease activity, we performed a fluorometric enzyme activity assay (Enzcheck) using the unselective serine protease trypsin (ThermoFisher Scientific, Waltham, MA, USA) at a concentration of 0.05%. Enzyme substrate was diluted in provided assay buffer according to manufacturer’s instruction. Equal amounts of Trypsin were 1:2 diluted in assay buffer, control medium or PBMCsec concentrated at 12.5 units/ml for five minutes before adding 10 μl to the prepared substrate mixture adding up to a total volume of 100μl per well. Urokinase inhibitor Screening Kit (Sigma-Aldrich, St. Louis, USA) was used to test the effect of the investigated plasma samples and PBMCsec on urokinse activity. In brief, 45 μl plasma sample were diluted in equal volume of provided assay buffer. Human urokinase and substrate were added as suggested by the protocol adding up to a total reaction volume of 100 μl per well. Samples from 3 donors were analyzed. For both tests, samples were then incubated at room temperature for a total of 60 minutes. Absorbance at 450 nm was measured by FluoStar Optima microplate reader (BMG Labtech, Ortenberg, Germany) in 15-minutes intervals.

### Tube formation assay

Pro-angiogenic properties of PBMCsec and plasma of PBMCsec-treated whole blood were compared in a tube formation assay with human umbilical vein endothelial cells (HUVECs, passage 8) as described previously [57]. After isolation, cells were routinely cultured in endothelial cell growth basal medium-2 (EBM-2; Lonza Group AG, Basel, Switzerland) supplemented with endothelial cell growth medium-2 (EGM-2; BulletKit, Lonza) until fully confluent. Prior to the tube formation assay, cells were maintained in EBM-2 containing 3% (vol/vol) heat-inactivated fetal bovine serum (Lonza) overnight and starved in basal EBM-2 without supplements for 3 hours. Cells were seeded on growth factor-reduced Matrigel Matrix (Corning Inc. Life Sciences, Tewksbury, MA, USA) in μ-slides Angiogenesis (ibidi GmbH, Graefelfing, Germany) at a density of 10 × 10^4^ cells per well and stimulated with the supernatant obtained from 3 ml whole blood cells for 4 hours. Micrographs were acquired by an inverted phase contrast microscope (CKX41 Olympus Corporation; Tokyo, Japan) equipped with a 10x objective (CAch N, 10x/0.25 PhP; Olympus) using a SC30 camera (Olympus) and cellSens Entry software (version 1.8; Olympus). Tubule formation was quantified by the Angiogenesis Analyzer plugin [58] of ImageJ (version 1.53, java 1.8.0_172) using default settings.

### Dermal Microvascular endothelial cell culture

Dermal microvascular endothelial cells (DMEC) were isolated from human foreskin. Foreskin was digested with dispase (Corning). Epidermis was removed and foreskin was scraped to dislodge endothelial cells. Cells were sorted for CD31 with magnetic beads (Thermo Fisher Scientific). Endothelial cells were cultured in endothelial growth medium (EGM-2; Lonza) containing 15% fetal bovine serum (FCSThermo Fisher Scientific, Waltham, MA, USA) and supplements for microvascular cells (Lonza). Cells were maintained in a humidified atmosphere containing 5% CO2 at 37°C and passaged at 90% confluence. Prior to experiments, cells were authenticated and confirmed to be free of contamination by mycoplasma. Endothelial cells were used at passages 2 to 8.

### Electrical cell-substrate impedance sensing (ECIS)

Electrical cell-substrate impedance sensing (ECIS, Applied Biophysics, Troy, NY, USA) was used to measure the electrical resistance of endothelial monolayers. 12 000 endothelial cells were seeded on array plates (Ibidi) coated with gelatin (Sigma). After resistance at 4000 Hz reached a stable plateau of >1000 Ω, endothelial cells were treated with indicated substances. Electrical resistance of cell monolayers was continuously monitored at 250 Hz [34].

### Statistical analysis

For single cell RNA seq, two donors were analyzed. Negative binomial regression was performed to normalize data and achieve variance stabilization. Wilcoxon rank sum test was followed by Bonferroni post-hoc test to calculate differentially expressed genes. For *in vitro* experiments, at least three different donors were used. For data analysis of tube formation assay, investigators were blinded to treatments. Data were statistically evaluated using GraphPad Prism v8.0.1 software (GraphPad Software, San Diego, USA). When analyzing three or more groups ordinary one-way ANOVA and multiple comparison post hoc tests with Dunnett’s correction were calculated and p-values < 0.05 were considered statistically significant. Data are presented as mean ± standard error of the mean (SEM).

### Patents

The Medical University of Vienna has claimed financial interest. HJA holds patents related to this work (WO 2010/079086 A1; WO 2010/070105 A1; EP 3502692; European Patent Office application # 19165340.1).

## Supporting information

supplementary sheet 1

supplementary sheet 2

## Author Contributions

M.M., H.J.A., and D.C. conceived and planned the experiments. H.J.A. and M.M. acquired funding. D.C., M.D., and K.S. performed the experiments. D.C., M.D., K.S., M.L., D.B., V.V., K.H., H.J.A. and M.M. participated in data interpretations. D.C., M.L. and M.M. drafted the manuscript. All authors reviewed the manuscript.

## Data Availability Statement

ScRNA-seq data are available upon request.

## Acknowledgments

We are thankful to Dr. HP Haselsteiner and the CRISCAR Familienstiftung for their belief in this private public partnership to augment basic and translational clinical research.

## Supplemental Information

**Supplementary Figure 1.**
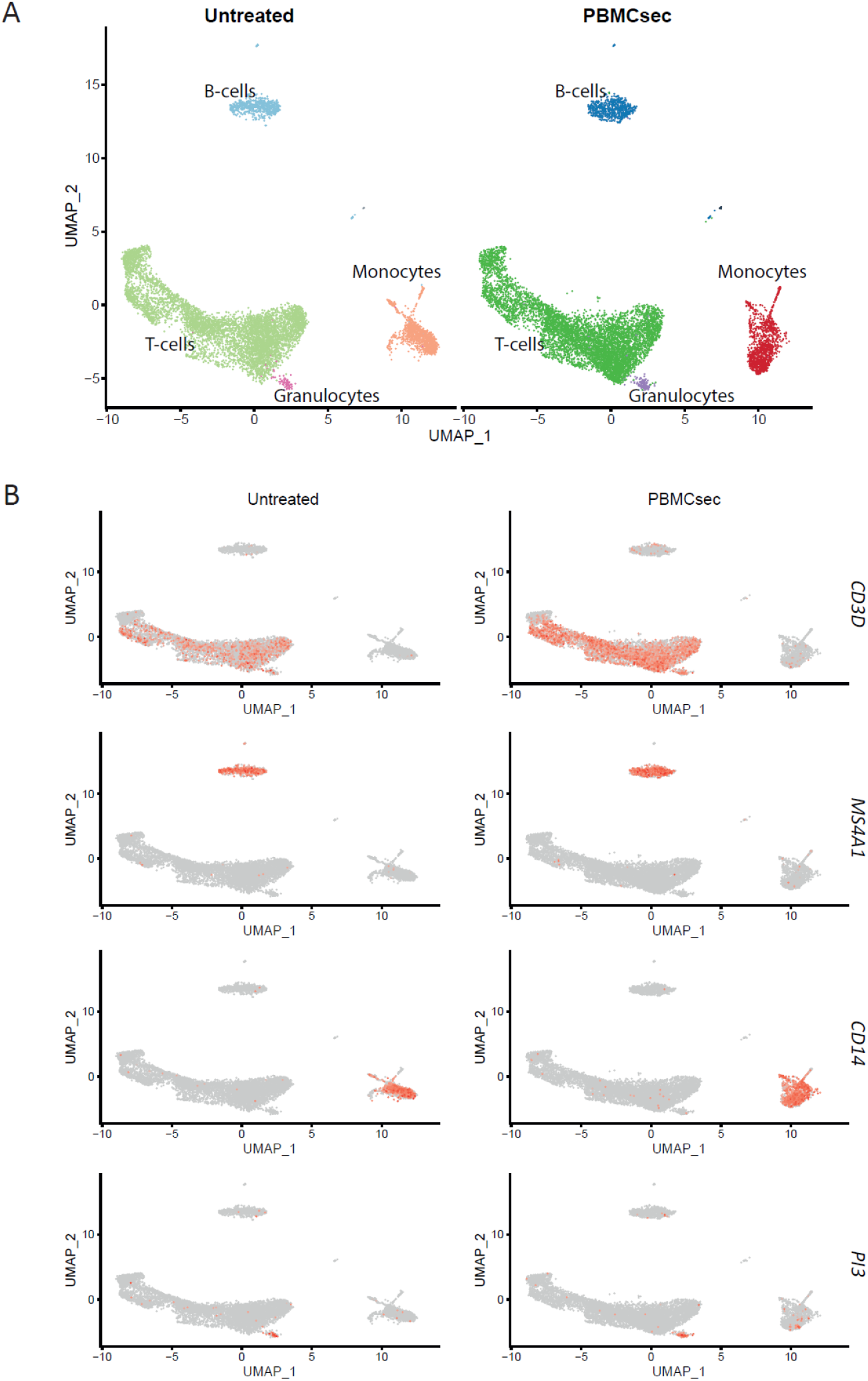
Identification of cell clusters based on expression of cluster-defining marker genes for T-cells, B-cells, Monocytes and granulocytes. (A) UMAP Plot with cell clusters and (B) Featureplots showing the average expression for *CD3D, MS4A1, CD14* and *PI3* in every cluster. Positive events for the respective marker gene are highlighted in red. Identified clusters are annotated as T-cells, B-cells, monocytes and granulocytes.

**Supplementary Figure 2.**
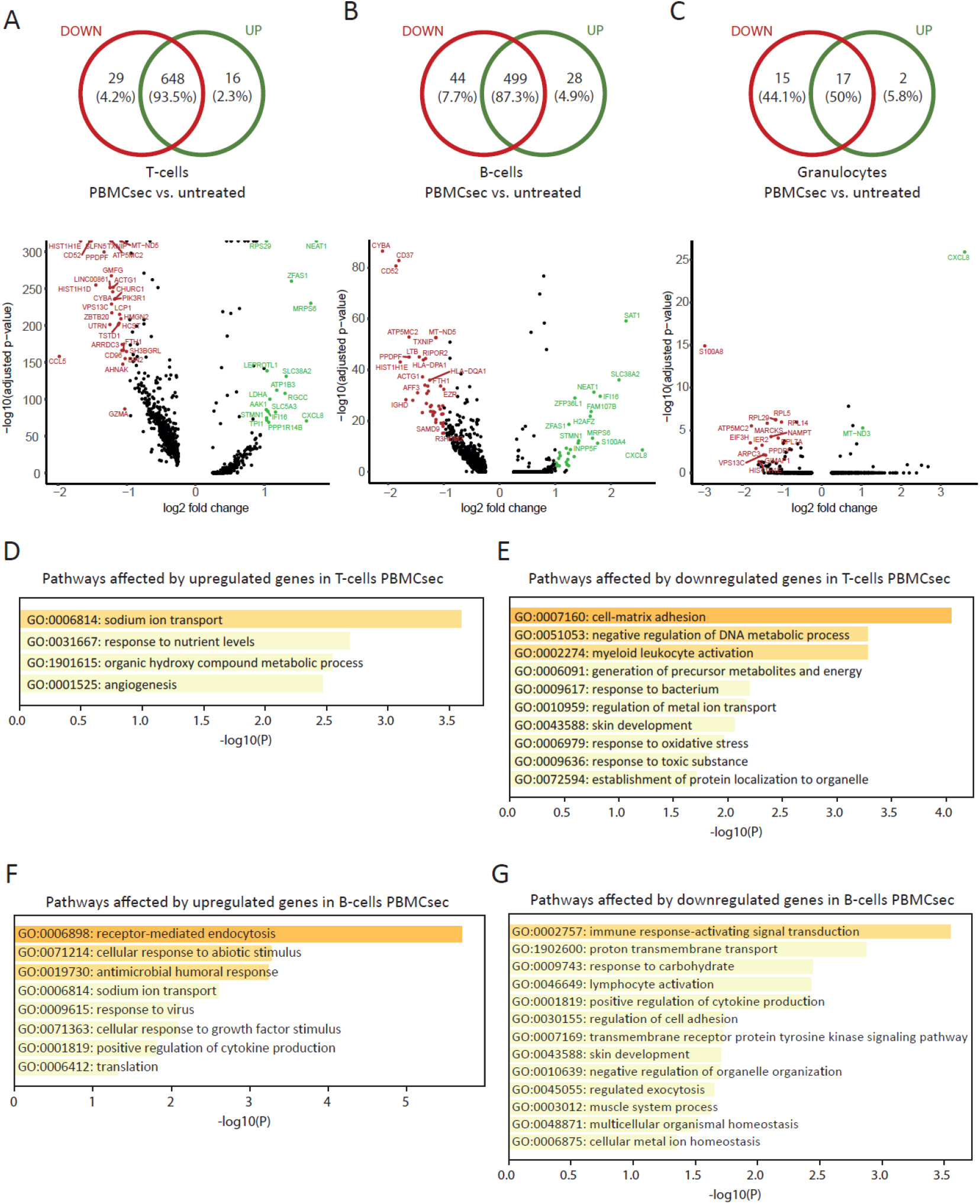
Transcriptional changes in T-cells, B-cells and granulocytes and their predicted regulation of biological processes. Distribution of up- and downregulated genes are shown for (A) T-cells, (B) B-cells and (C) granulocytes. Barplots display the significantly enriched biological processes affected by (D) upregulated and (E) downregulated gene sets for T-cells. Barplots display the significantly enriched biological processes affected by (F) upregulated and (G) downregulated gene sets for B-cells.

**Supplementary Figure 3.**
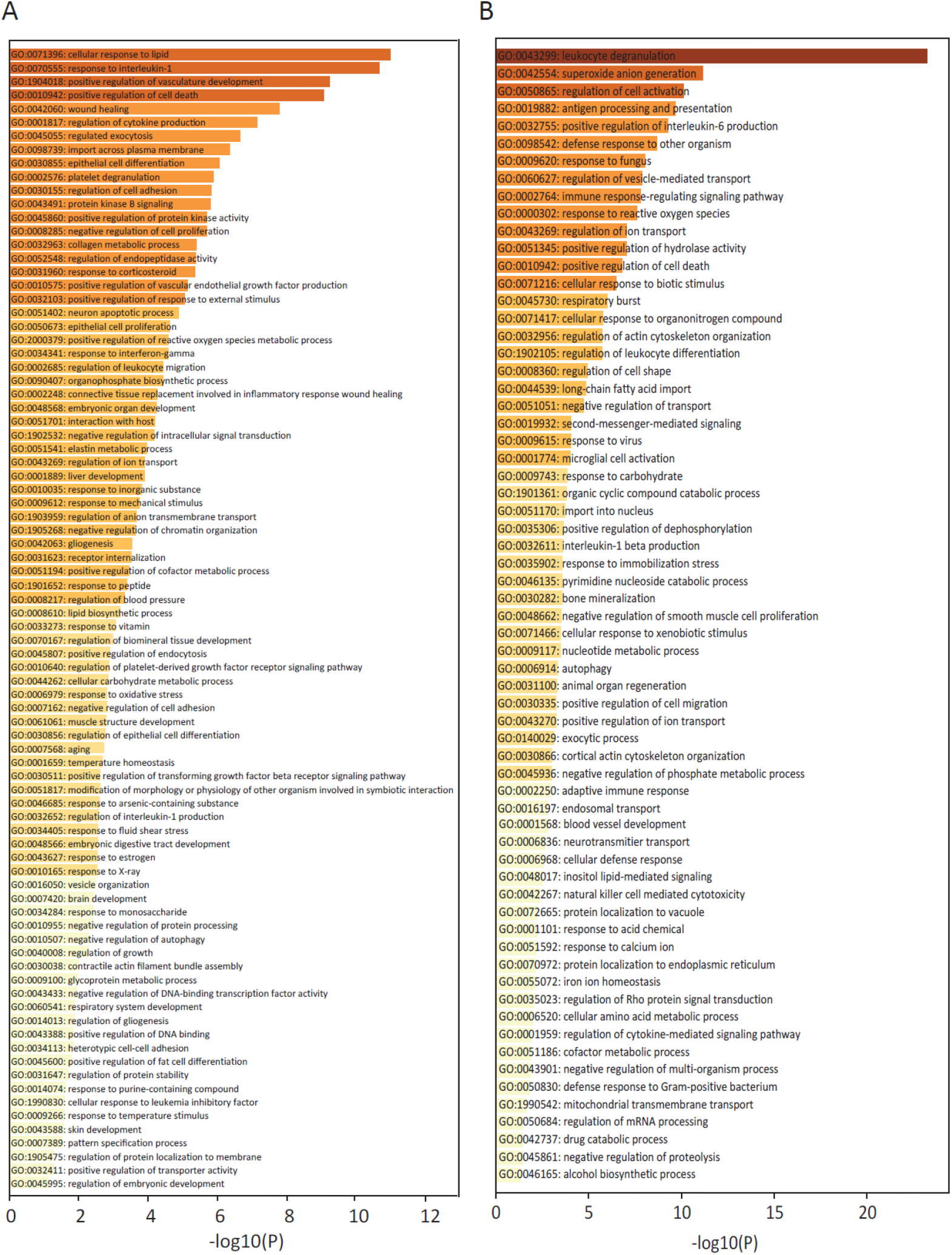
Biological processes affected by up- and downregulated genes in monocytes treated with PBMCsec. (A) Barplot of significantly biological processes enriched by the the gene set of upregulated genes and (B) downregulated genes in monocytes treated with PBMCsec.

**Supplementary Figure 4.**
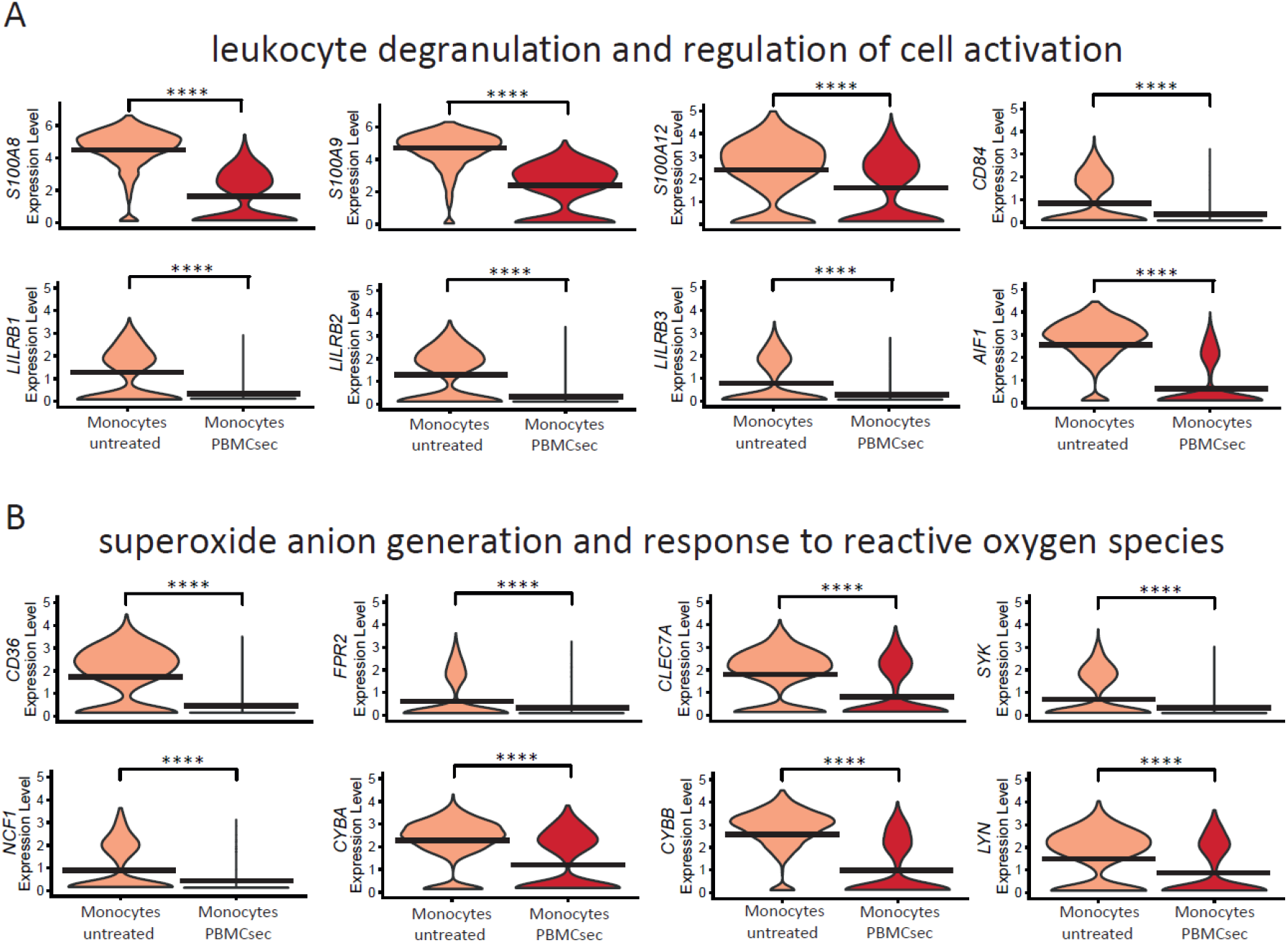
Downregulated genes in monocytes treated with PBMCsec associate with leukocyte degranulation and generation of reactive oxygen species. (A) Violin Plots show a selection of most downregulated genes that contribute to leukocyte degranulation and activation and (D) superoxide anion generation and response to reactive oxygen species. **** indicate p-value < 0.0001.

**Supplementary sheet 1**. Pathways associated with significantly upregulated genes in monocytes treated with PBMCsec.

**Supplementary sheet 2**. Pathways associated with significantly downregulated genes in monocytes treated with PBMCsec.

## Graphical summary

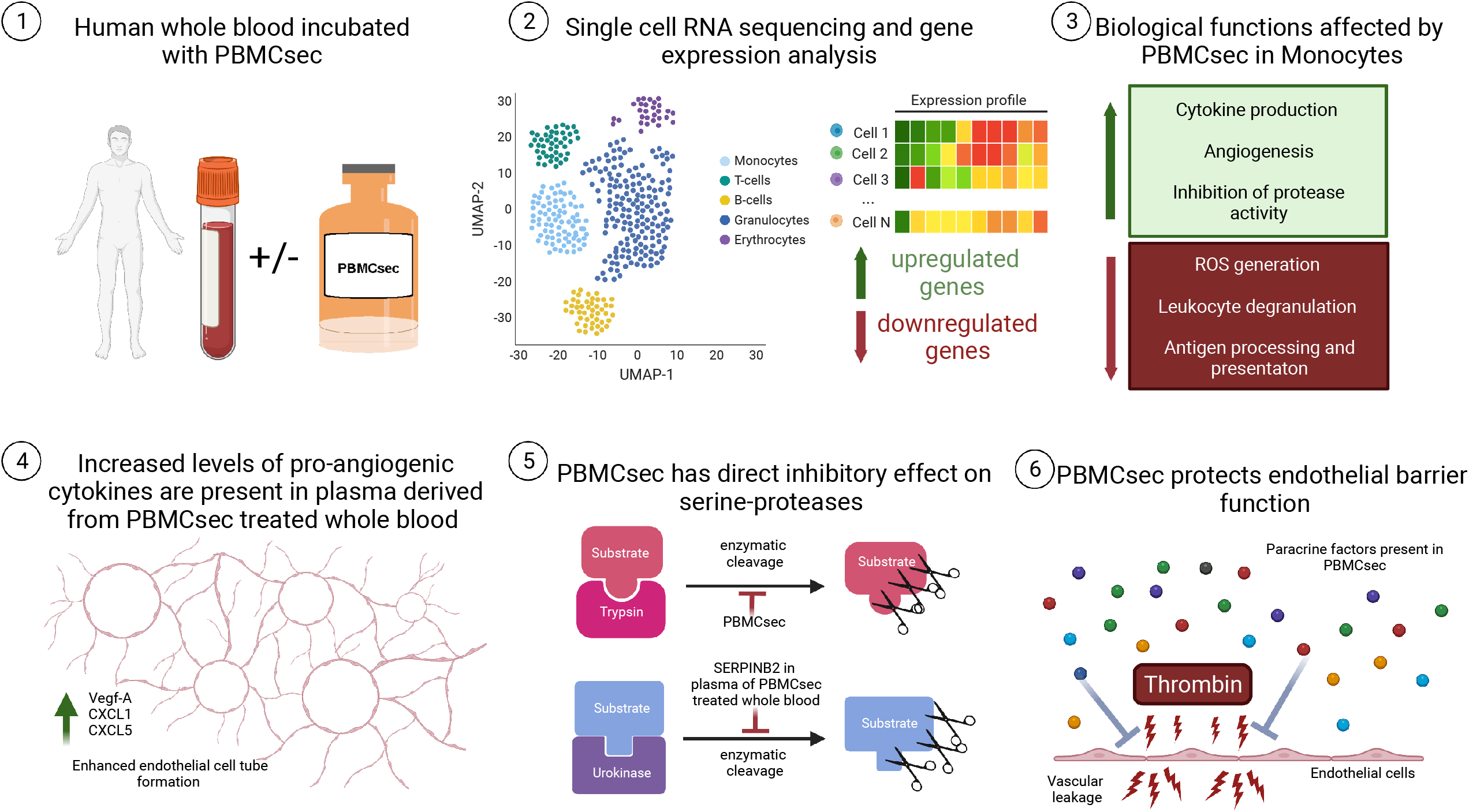

